# Spinal cord and brain concentrations of riluzole after oral and intrathecal administration: a potential new treatment route for amyotrophic lateral sclerosis

**DOI:** 10.1101/2022.11.02.514962

**Authors:** Orion P. Keifer, Juan Marco Gutierrez, Mark T. Butt, Sarah D. Cramer, Raymond Bartus, Malu Tansey, Daniel Deaver, Alexandre Betourne, Nicholas M Boulis

## Abstract

Riluzole is the only treatment known to improve survival in patients with Amyotrophic Lateral Sclerosis (ALS). However, oral riluzole efficacy is modest at best, further it is known to have large inter-individual variability of serum concentration and clearance, is formulated as an oral drug in a patient population plagued with dysphagia, and has known systemic side-effects like asthenia (limiting patient compliance) and elevated liver enzymes. In this context, we postulated that continuous intrathecal (IT) infusion of low doses of riluzole could provide consistent elevations of the drug spinal cord (SC) concentrations at or above those achieved with oral dosing, without increasing the risk for adverse events associated with systemic drug exposure or off-target side effects in the brain. We developed a formulation of riluzole for IT delivery and conducted our studies in purpose-bred hound dogs. Our non-GLP studies revealed that IT infusion alone was able to increase SC concentrations above those provided by oral administration, without increasing plasma concentrations. We then conducted two GLP studies that combined IT infusion with oral administration at human equivalent dose, to evaluate SC and brain concentrations of riluzole along with assessments of safety and tolerability. In the 6-week study, the highest IT dose (0.2 mg/hr) was well tolerated by the animals and increased SC concentrations above those achieved with oral riluzole alone, without increasing brain concentrations. In the 6-month study, the highest dose tested (0.4 mg/hr) was not tolerated and yielded SC significantly above those achieved in all previous studies. Our data show the feasibility and safety profile of continuous IT riluzole delivery to the spinal cord, without concurrent elevated liver enzymes, and minimal brain concentrations creating another potential therapeutic route of delivery to be used in isolation or in combination with other therapeutics.”

## Introduction

Amyotrophic lateral sclerosis (ALS) is a rapidly progressive, neurodegenerative disease that affects upper and lower motor neurons in the brain and spinal cord (SC) [1]. It is the most common motor neuron disorder, with a prevalence of approximately 5 per 100,000 [2]. ALS results in irreversible wasting and paralysis of voluntary and involuntary muscles and motor function [3, 4]. Though many different phenotypes of ALS exist, the relatively rapid progression and primary neuromuscular symptoms are shared among the vast majority of cases. All patients get weaker and less mobile over time, which substantially reduces the patients’ quality of life, and often requires feeding tubes due to the inability to swallow, and non-invasive respirators to aid breathing [5]. While the rate of progression from symptom onset to death can vary significantly among patients, the typical disease duration is 2.5-5 years and approximately 50% of ALS cases dying within three years most often due to diaphragmatic failure [6-10].

No current treatment prevents, halts, or reverses the disease. Two treatments for ALS have been approved by the FDA: edavarone and riluzole. Edavarone was approved by the FDA in 2017 but has not yet been shown to extend patient survival [11, 12]. Riluzole (50 mg orally, twice daily) has only very modest efficacy, providing a survival benefit of approximately 3 months [13, 14]. While riluzole’s mechanism of action is not fully understood, it is known to be a sodium channel blocker, which reduces intracellular sodium leading to less neuronal membrane depolarization [15, 16]. Riluzole also reduces glutamate release and increases synaptic glutamate uptake, thereby reducing NMDA receptor-mediated neuronal excitotoxicity [17]. It is through either or both of these mechanisms that riluzole is believed to exert its neuroprotective effects on the motor neurons [17]. Although ALS affects both upper and lower motor neurons, evidence supports the hypothesis that part of the benefit of riluzole may be achieved through its action on lower motor neurons in the SC.

Unfortunately, not all patients tolerate riluzole, as evidenced by a dose dependent increase in patient adverse events and withdrawal during dose ranging studies. Particularly notable are dose-dependent increases in asthenia and digestive issues which limit patient compliance (see [14, 18]). Further, there is also a dose-related effect on liver enzymes, particularly the time to liver enzyme rise (earlier in higher doses) and increases with ALT ≥ 3 x upper limit of normal (ULN) but < 5 ULN – though, interestingly, > than 5x ULN, the criteria for withdrawal, was not dose dependent [14, 18].In a clinical setting, these elevations resolve at the currently accepted 50 mg BID dose, though it is noted that special care and caution is advised with patients with current or past abnormal liver function (Rilutek® package insert, 2016 [50]). Additionally, there is significant inter-patient variability with the clearance and thus plasma concentrations of the riluzule which makes it difficult to consistently dose patients to a known therapeutic value (i.e.100 mg for one patient does not equal the same therapeutic dose on the spinal cord and brain) [19]. Further, riluzole is a known substrate of the p-glycoprotein system, thus limiting its blood-brain barrier penetration (augmenting the challenges of assuring appropriate therapeutic dose on target), which could be further exacerbated by evidence showing that p-glycoprotein system is upregulated in ALS [20-22]. Finally, but not insignificantly, current formulation of riluzole are oral, which is particularly challenging in a patient population plagued with high incidences of dysphagia [23]. Crushing the pills complicates the already unpredictable pharmacokinetics and increases the variability of dosing [23, 24]. Continuous intrathecal delivery delivery of riluzole using an implanted pump has the potential to ameliorate many of these concerns, and also potentially increase the concentration of riluzole in the spinal cord beyond what is accomplished with the current oral dosing.

The current oral dose for riluzole is 100mg, based on clinical trials showing little benefit of the 200mg dose, while showing increases in side effects including asthenia elevated ALT levels, which were “more common in the higher (i.e., 200 mg/day) dose” (NDA 20-599). These observations led the investigators to conclude that with orally-administered riluzole, “overall, efficacy and safety results suggest that the 100 mg [daily])dose has the best benefit-to-risk ratio” [14]. To that point, it has been noted that “the modest efficacy of riluzole on [ALS] disease progression may be the price to pay for acceptable safety of the drug” [25]. Yet, evidence exists that doses higher than 100 mg per day could potentially be more effective on the desired target tissue. For example, a number of investigators have tested higher riluzole doses (up to 200 mg daily) for a variety of CNS indications (both neurological diseases and psychiatric disorders) and reported positive effects at daily doses above 100 mg [26-30]. These data indicate that higher SC levels might provide greater benefit to ALS patients.

For these reasons, we hypothesized that we could establish consistent SC concentrations of riluzole, at or above that available from oral dosing, with local delivery in the cerebrospinal fluid (CSF). Further, we hypothesized that we could also mitigate the ALT effects present with oral riluzole. In order to empirically test this concept, a riluzole formulation was developed to enable riluzole delivery through an intrathecal pump and four separate studies were conducted. First, a non-GLP study was performed to evaluate the pharmacokinetics of IT riluzole in comparison with the oral route of administration. Then, three studies were conducted to evaluate the safety and tolerability of IT administered riluzole in purpose-bred hound dogs. Importantly, incorporated into each study was a subset of experiments focused on the tissue distribution of riluzole to SC, brain, and systemic blood. The first of these studies was a non-GLP dose-escalation study using IT riluzole only, which allowed for an assessment of SC concentrations of IT riluzole alone. This was followed by 6-week and 6-month GLP studies that combined IT infusion and oral riluzole to evaluate SC and brain concentrations of riluzole achieved with the combination along with assessments of safety and tolerability. The combination of oral and IT riluzole was evaluated as it is unlikely that patients would discontinue oral riluzole during assessment of the safety and efficacy of intrathecal riluzole, thus these studies were completed to enable inclusion in an IND-enabling package. This publication focuses on SC, brain, and systemic blood plasma data, but information on overall clinical tolerability and local histopathology associated with IT infusion are also summarized.

We propose that IT-riluzole, either with or without concurrent oral riluzole, may represent a feasible and effective means of enhancing riluzole concentration to the spinal cord. Further evaluation of the efficacy of the treatment in a prospective fashion is needed.

## Methods

### Animals

All studies were approved by the Institutional Animal Care & Use Committee (IACUC) at MPI Research and conducted within MPI facilities (now Charles River Laboratories, Inc. Mattawan, MI). Animal Welfare Standards followed U.S. Department of Agriculture’s (USDA) Animal Welfare Act (9 Code of Federal Regulations (CFR) Parts 1, 2, and 3) and Guide for the Care and Use of Laboratory Animals. 8th ed., Washington, D.C., National Academies Press, 2011. Purpose-bred hounds were acquired from Marshall Bioresources (North Rose, NY) with a weight of 20-30 kg. Prior to study start animals were group housed. Following surgery, animals were individually housed in single- or double-sized cages. The dogs were provided the opportunity for exercise according to MPI Research internal Standard Operating Procedures (SOP). The caging used runs with raised flooring, a type of housing considered to provide adequate room to exercise for these animals. Temperature and humidity were maintained according to MPI Research internal SOP. Fluorescent lighting was provided via an automatic timer for approximately 12 hours per day. On occasion, the dark cycle may have been interrupted intermittently due to study-related activities. Tap water was supplied ad libitum to all animals via an automatic water system unless otherwise indicated. Lab Diet® Certified Canine Diet #5007, PMI Nutrition International, Inc. was provided ad libitum to all animals at arrival, except during fasting periods or select husbandry functions (e.g., exercise and room cleaning). Animal enrichment was provided according to MPI Research internal SOP. Cageside observations were performed at least twice daily and animals were observed for morbidity, mortality, injury, and availability of food and water.”

### Surgical Procedures

The Medtronic AscendaTM intrathecal catheter (model 8780) and Medtronic Synchromed® II pump (model 8637-40ml reservoir) were selected to deliver the test article into the subdural space, due to ease of programming multiple infusion rates and its prior use in human ALS patients [31]. The purpose-bred hound dog was selected for these studies over alternatives such as pure-bred beagles or pigs because their larger size better accommodates the human intrathecal pump without skin breakdown. Furthermore, the purpose-bred hound dog vertebral column and SC anatomical features are more similar to the human and thus better suited to accommodate the human catheter size and representatively model SC distribution achieved during IT dosing.

Intrathecal catheter placement is commonly performed via percutaneous insertion in humans. However, pilot experiments showed that this approach was associated with unacceptable morbidity and frequent mispositioning of the catheters when performed in dogs. As such, open placement was employed for dog studies, and a two-level laminectomy was performed to implant the Ascenda catheter. Surgeries were performed at least 10 days before dosing began to allow time for the animals to recover. At surgery, anesthetized dogs were scrubbed with chlorhexidine and draped for surgery. A 10cm incision was made to expose the lamina, and then a two-level laminectomy was performed in the lumbar spine. The catheter was introduced into the subarachnoid space with a curved Tuohy needle that was passed through the paraspinous muscle caudal to the laminectomy under direct visualization. A 4-0 stitch was used to elevate and immobilize the dura, limiting damage to underlying nerve roots. The catheter was advanced under fluoroscopic guidance (to the thoracic vertebrae T4 in Experiment 2, to the cervicothoracic junction in Experiments 1, 3 and 4). Catheter location in the subdural space was confirmed with fluoroscopy and myelography (Fig 1). A 4-0 purse string stitch was placed around the catheter at the dura, and a second 2-0 purse-string suture was used to minimize leakage of cerebrospinal fluid into the subcutaneous space. The IT catheter was tunneled through muscle and subcutaneous tissue, where it was anchored with a strain relief loop in a subdermal pocket formed to accommodate the pump implantation in the animal’s flank through a separate incision. The pump was wrapped in Dacron felt (promote scaring to help limit seroma formation) and secured to the fascia and interspinous ligaments with sutures to limit migration. At the time of implantation, the pump was filled with saline and set to the minimum flow rate to ensure catheter patency (0.006 mL/day). The incisions were closed in layers in a standard fashion.

**Fig 1.**
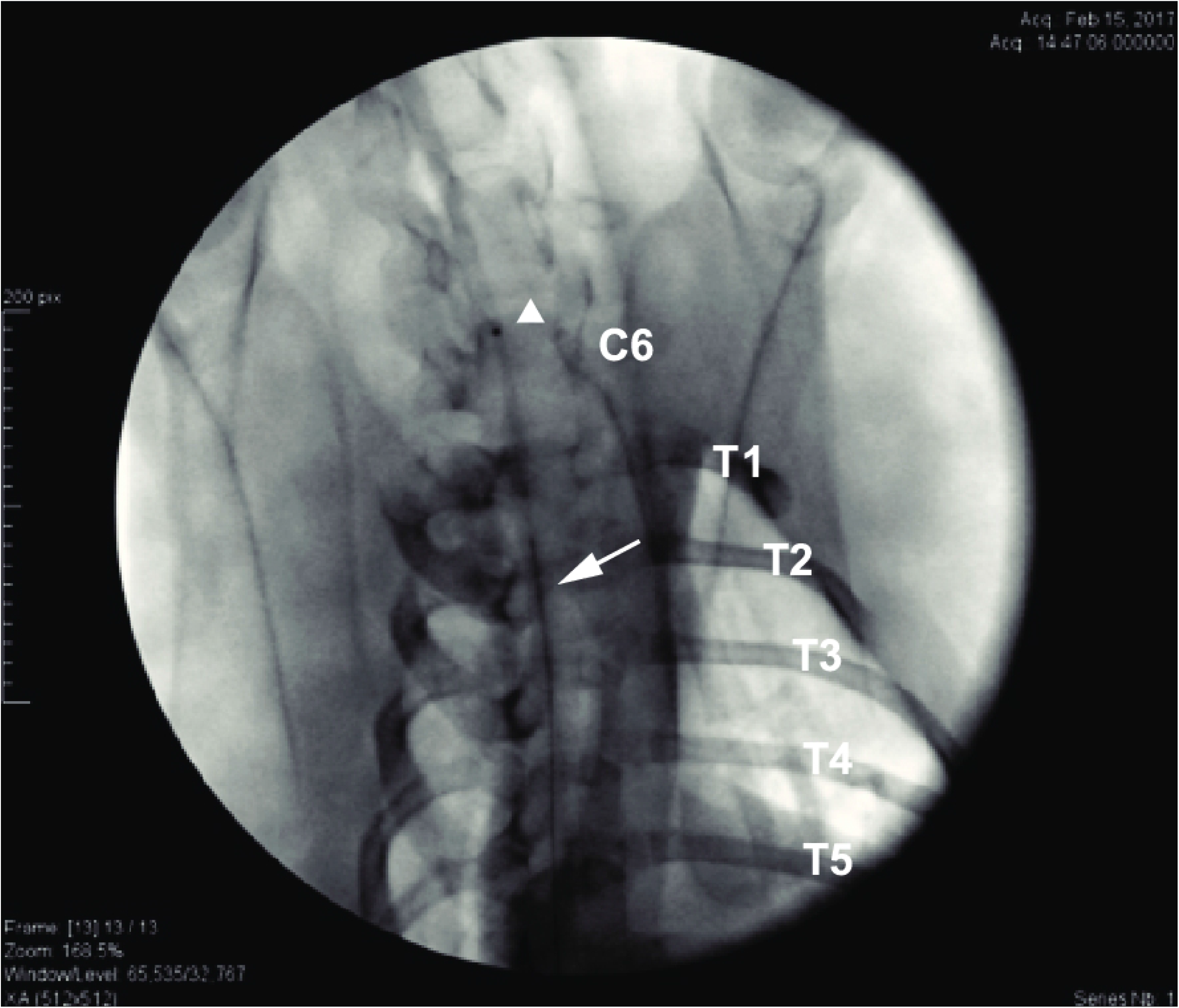
Myelogram obtained during catheter implant at surgery. The radiopaque catheter tip guides the positioning at the cervico-thoracic junction. Contrast agent is flushed through the pump access port to confirm correct positioning in the subdural space. White arrow points to the catheter and white arrowhead to the catheter tip. C6: cervical vertebrae 6, T1-5: thoracic vertebrae 1-5

### Drug Preparations of Riluzole

Riluzole was administered orally by powder (Riluzole cGMP API, ScinoPharma, Taiwan) compounded in #12 gelatin capsules (Torpac, Fairfield, NJ) (Study 2) or commercially available USP tablets (Medline Industries Inc., Libertyville, Illinois) (Studies 1, 3 and 4). For IT administration, riluzole was formulated in Trappsol Cyclo or Trappsol® Hydroxypropyl Beta Cyclodextrin Endotoxin Controlled (THPB-EC; CTD Holdings, Inc., Alachua, FL) to increase the drug solubility. For Study 1, a 4 mg/mL formulation of riluzole was prepared in 4% THPB-EC in a sodium chloride/sodium phosphate solution prepared in Sterile water for injection (USP, pH 7.4), and pH was adjusted to neutrality. For Study 2, riluzole was diluted in 5% Trappsol Cyclo in 0.9% Sodium Chloride for Injection, USP (B. Braun Medical Inc., Bethlehem, PA) to produce the following doses: 1.82 mg/mL, 2.73 mg/mL, 3.64 mg/mL, 4.55 mg/mL. For Study 3, a 4 mg/mL formulation of riluzole was prepared in 4% THPB-EC in a sodium chloride/sodium phosphate solution prepared in Sterile water for injection (USP, pH 7.4), and pH was adjusted to neutrality. The mid and low IT doses (2 mg/ml and 1 mg/ml, respectively) were then prepared by serial dilution in 0.9% saline (USP). For Study 4, the riluzole formulation preparation was identical to Study 3, but the low IT dose (1mg/ml) was prepared by dilution in the THPB-EC vehicle: 4% THPB-EC in a sodium chloride/sodium phosphate solution prepared in sterile water for injection. See Table 1 for riluzole formulations summary.

**Table 1.**
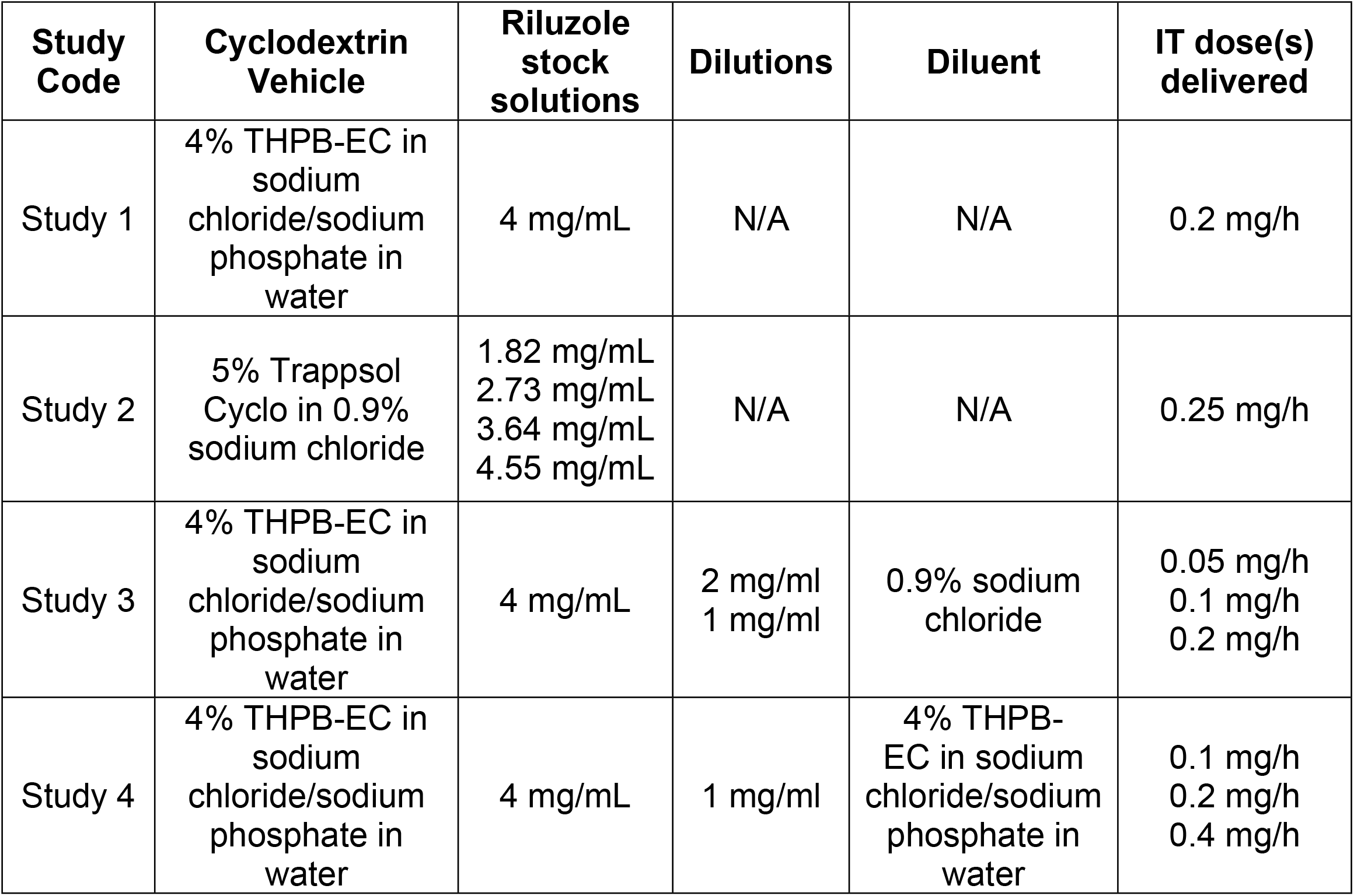
Summary of riluzole IT formulations.

| **Study Code** | **Cyclodextrin Vehicle** | **Riluzole stock solutions** | **Dilutions** | **Diluent** | **IT dose(s) delivered** |
| --- | --- | --- | --- | --- | --- |
| Study 1 | 4% THPB-EC in sodium chloride/sodium phosphate in water | 4 mg/mL | N/A | N/A | 0.2 mg/h |
| Study 2 | 5% Trappsol Cyclo in 0.9% sodium chloride | 1.82 mg/mL<br>2.73 mg/mL<br>3.64 mg/mL<br>4.55 mg/mL | N/A | N/A | 0.25 mg/h |
| Study 3 | 4% THPB-EC in sodium chloride/sodium phosphate in water | 4 mg/mL | 2 mg/ml<br>1 mg/ml | 0.9% sodium chloride | 0.05 mg/h<br>0.1 mg/h<br>0.2 mg/h |
| Study 4 | 4% THPB-EC in sodium chloride/sodium phosphate in water | 4 mg/mL | 1 mg/ml | 4% THPB-EC in sodium chloride/sodium phosphate in water | 0.1 mg/h<br>0.2 mg/h<br>0.4 mg/h |

### Study Designs

The studies of IT riluzole in purpose-bred hound dogs are summarized in Table 2.

**Table 2.** Studies summary.

| **Study Code** | **GLP Status** | **Study type** | **Study Title** |
| --- | --- | --- | --- |
| Study 1 | Non-GLP | PK | Pharmacokinetic study of IT riluzole |
| Study 2 | Non-GLP | Safety/PK | Five-day Dose Escalation Study Using IT Infusion of Riluzole Only |
| Study 3 | GLP | Safety/Toxicology | GLP 6-week Safety and Tolerability Study Using the Combination of Oral and IT infusion of Riluzole |
| Study 4 | GLP | Safety/Toxicology | GLP 6-Month Safety and Tolerability Study Using the Combination of Oral and IT infusion of Riluzole |

#### Study 1: Pharmacokinetic study of IT riluzole

Six purpose-bred male hound dogs were used for this study. Three dogs were assigned to receive continuous IT infusion of riluzole at 0.2 mg/hr for 5 days. On the first day of dosing, the pump reservoir containing saline was emptied then filled with 40mL of drug solution. A priming bolus based on the length of the catheter was programmed to bring the drug solution to the tip of the catheter. IT infusion started thereafter at the programmed rate of 1.2 mL per day to deliver a riluzole dose of 0.2 mg/hr. Plasma samples were taken from the jugular vein at 0.5, 1, 1.5, 2, 4, 8, 12, 24, 32, 48, 56, 72, 80, 96, 104, 120, and 128 hours from the start of infusion.

The three other dogs received a single 50 mg dose of riluzole administered orally, followed by a tap water rinse. Plasma samples were taken from the jugular vein at 0.25, 0.5, 1, 1.5, 2, 4, 8, 12, 24, 48, and 72 hours post-dose.

#### Study 2: Five-day Dose Escalation Study Using IT Infusion of Riluzole Only

Ten purpose-bred hound dogs were used for this study. Six dogs (three males and three females) were assigned to receive four IT doses of riluzole (0.10, 0.15, 0.20 and 0.25 mg/hr) in a dose-ascending order; all six dogs received all four doses. On the first day of dosing, the pump reservoir containing saline was emptied and filled with drug solution. The infusion rate during each 5-day treatment period was 1.32 mL/day, and was switched to the pump minimum rate (0.006 mL/day) to ensure catheter patency during the two days interval between each dose, during which the pump reservoir was flushed and re-filled with the next dose. At the end of the last infusion period (0.25 mg/hr dose), dogs were euthanized to collect SC and brain tissues for analysis of riluzole concentrations.

To compare systemic concentrations of riluzole achieved with IT infusion to those achieved with oral dosing, 4 dogs (two males and two females) received oral riluzole at HED. Riluzole was filled into individual capsules to achieve a dose of 50 mg given twice daily (BID) for 14 days. At the end of the 14-day treatment period, blood was obtained approximately 1 hour after the last oral dose (estimated time of C_max_ from literature), and dogs were euthanized to collect SC and brain tissues for analysis of riluzole concentrations. Because no differences between males and females were observed, bioanalytical data were pooled for presentation.

#### Study 3: GLP 6-week Safety and Tolerability Study Using the Combination of Oral and IT infusion of Riluzole

Thirty-six male purpose-bred hound dogs were used for this Food and Drug Administration (FDA) Good Laboratory Practices (GLP)-compliant study. Dogs were randomly assigned to one of six treatment groups: 1) IT infusion of saline; 2) IT infusion of drug vehicle; 3) oral treatment only with riluzole 50 mg BID (note: this group did not receive any surgery); 4) oral riluzole 50 mg BID (i.e. HED) and 0.05 mg/hr IT riluzole; 5) oral riluzole 50 mg BID and 0.1 mg/hr IT riluzole; and 6) oral riluzole 50 mg BID and 0.2 mg/hr IT riluzole. On the first day of dosing, the pump reservoir containing saline was emptied and filled with drug solution. The pump programmed infusion rate was then switched from minimum rate to 1.2 mL per day in all IT groups, to perform continuous infusion for 6-weeks (with a pump refill at 3-weeks). At the end of the in-life phase, dogs (n=6 per group) were euthanized to collect SC and brain tissues for analysis of riluzole concentrations and histopathology. In addition, blood and peripheral organs/tissues were collected for standard clinical chemistry and histopathology toxicology assessments performed according to MPI Research SOP. For sake of brevity, we only present the hepatic function assessment, evaluated by measuring the activities of the liver enzymes alanine aminotransferase (ALT), aspartate aminotransferase (AST) and gamma-glutamyltransferase (GGT).

#### Study 4: GLP 6-Month Safety and Tolerability Study Using the Combination of Oral and IT infusion of Riluzole

Thirty-nine male purpose-bred hound dogs were used for this FDA GLP-compliant study. Dogs were randomly assigned to one of six treatment groups: 1) IT infusion of saline; 2) IT infusion of drug vehicle; 3) oral treatment only with 50 mg riluzole BID (note: this group did not receive any surgery); 4) oral riluzole 50 mg BID and 0.1 mg/hr IT riluzole; 5) oral riluzole 50 mg BID and 0.2 mg/hr IT riluzole; and 6) oral riluzole 50 mg BID and 0.4 mg/hr IT riluzole. On the first day of dosing, the pump reservoir containing saline was emptied and filled with drug solution. The pump programmed infusion rate was then switched from minimum rate to 1.2 mL per day in group 4 and 5, and to 2.4 mL per day in group 6, to perform continuous infusion for 24-weeks (with monthly pump refills). At the end of the in-life phase, dogs (n=6-7 per group) were euthanized to collect SC and brain tissues for analysis of riluzole concentrations and histopathology. In addition, blood and peripheral organs/tissues were collected for standard clinical chemistry and histopathology toxicology assessments performed according to MPI Research SOP. For sake of brevity, we only present the activities of the liver enzymes ALT, AST and GGT. Several animals from group 6 were removed from the study early to determine spinal cord concentrations of riluzole at a time when clinical signs were present, and end of life collections were performed as described above.

### Drug Analysis

Riluzole was assayed by ultra-performance liquid chromatography with tandem mass spectrometry (UPLC with MS/MS) using positive ion electrospray, after isolation through supported liquid extraction at PPD labs (Middleton, Wisconsin). The assay was validated for both plasma and nervous tissue matrixes. The limit of quantitation was 0.500 ng/mL, and relative errors for accuracy and precision within the linearity range (0.5-500 ng/mL) were both less than ±10%.

### Necropsy

Following blood collection and fluoroscopic verification of catheter localization, animals were sedated with dexmedetomidine and received intravenous sodium heparin (200 IU/kg) followed by propofol (8mg/kg) then potassium chloride (1-2 mEq/kg). Animals were exsanguinated and then perfused with 0.9% saline at a rate of 0.5 to 1.5 L/min for 5 minutes. Animals were examined externally and internally throughout the abdominal, thoracic, and cranial cavities for any abnormalities. Tissues that were processed for histology were fixed in neutral buffered formalin for microscopic evaluation (Studies 2, 3, and 4).

For Studies 2, 3, and 4, brains were cut into 5 mm coronal sections and divided for histopathology and bioanalysis. After SC removal, the location of the catheter tip was noted using a tissue dye. For the SC, a 2.5cm section centered on the catheter tip location at necropsy was taken for histology, with a 3mm subsection (caudal and/or rostral to the catheter tip location) allotted for bioanalysis. For Studies 3 and 4, the remainder of the SC was also preserved, with alternating 1 cm sections used for histopathology and bioanalysis. For non-implanted animals (oral groups), the thoracic vertebrae T1 was considered to be the location of the catheter tip for anatomical orientation and continuity among groups. Spinal nerve roots and dorsal ganglia (DRG) from similar levels as the spinal cord sections were also collected for histopathology. Peripheral nerves, including the sciatic, tibial, and sural peripheral nerves, were collected and fixed in 2% methanol-free formaldehyde/2.5% glutaraldehyde then transferred to phosphate buffered saline.

For microscopic examination, fixed tissues were processed, embedded in paraffin, sectioned using a 5-micron block advance, and mounted to slide. Transverse sections of peripheral nerves were osmicated prior to paraffin-embedding. All slides were stained with hematoxylin and eosin (H&E). Slides of spinal cord were also labeled with ionized calcium binding adaptor molecule 1 (IBA-1) immunohistochemistry (IHC) for microglial cells and glial fibrillary acidic protein (GFAP) IHC for astrocyte reactions. Tissue histopathology was evaluated by board-certified veterinary pathologists (SDC, MTB).

### Pharmacokinetic Analysis

Study 1: for each animal, the following pharmacokinetic parameters were determined: maximum observed plasma concentration (C_max_) for the oral and IT infusion routes of administration, time of maximum observed plasma concentration (T_max_) for the oral and IT infusion routes of administration, and area under the plasma concentration-time curve (AUC). The AUC from time 0 to 72 hours (AUC0-72hr) for the oral route of administration, the AUC from time 0 to 128 hours (AUC0-128hr) for the IT infusion route of administration, the AUC from 48 hours to 128 hours for the IT route of administration, the AUC from time 0 to the time of the final quantifiable sample (AUCTlast), and the AUC from time 0 to infinity (AUCINF) for the oral route of administration were calculated by the linear trapezoidal method for all animals with at least three consecutive quantifiable concentrations. The relative BA of IT riluzole was calculated as follows: IT AUC (0-128 hr)/ Oral AUC(Inf)*100.

### Statistics

GraphPad Prism (Version 8) was used to create all data figures and, when applicable, for statistics, Data shown in Figs 2-5 represent means +/- SEM, data shown in Figs 6-7 represent means + SEM. A linear regression was calculated for Fig 3, and a correlation between plasma and SC concentrations observed in dogs receiving oral therapy only was plotted on Fig 6. When indicated, brain concentrations were analyzed using a one-way analysis of variance (ANOVA). Post hoc multiple comparisons were carried out when allowed, using Tukey’s Multiple Range test. Alpha levels were set at P < 0.05 for all tests.

**Fig 2.**
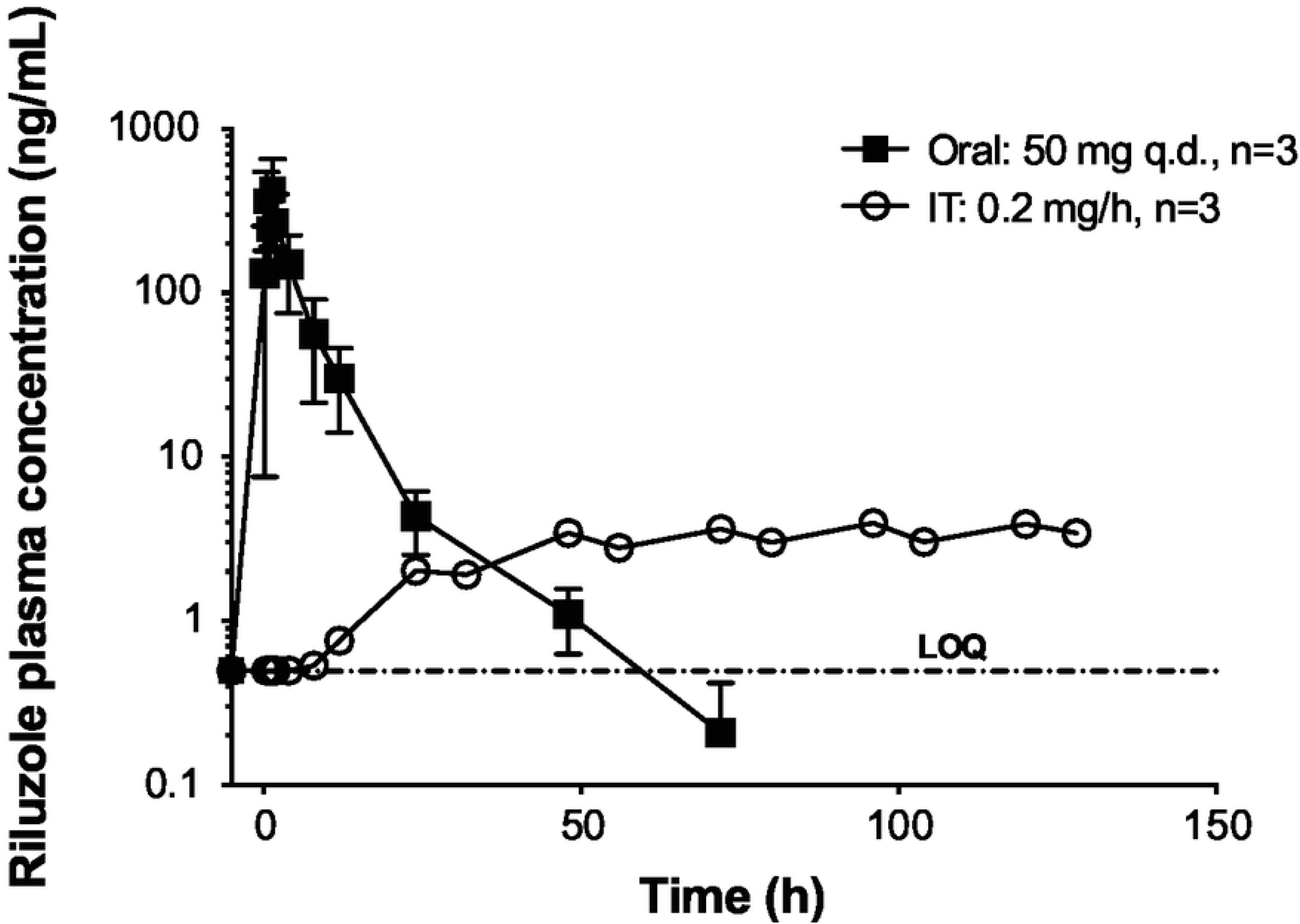
Study 1: Riluzole plasma concentrations. Riluzole plasma concentrations were measured after oral administration (50 mg, single dose) and during continous IT infusion of riluzole at 0.2 mg/h for 128 hours in dogs from study 1. Data shown as means ± SEM (some error bars are shorter than the symbols and cannot be visualized). Data below the assay limit of quantitation (BLOQ) are reported at the BLOQ value (0.500 ng/mL)

**Fig 3.**
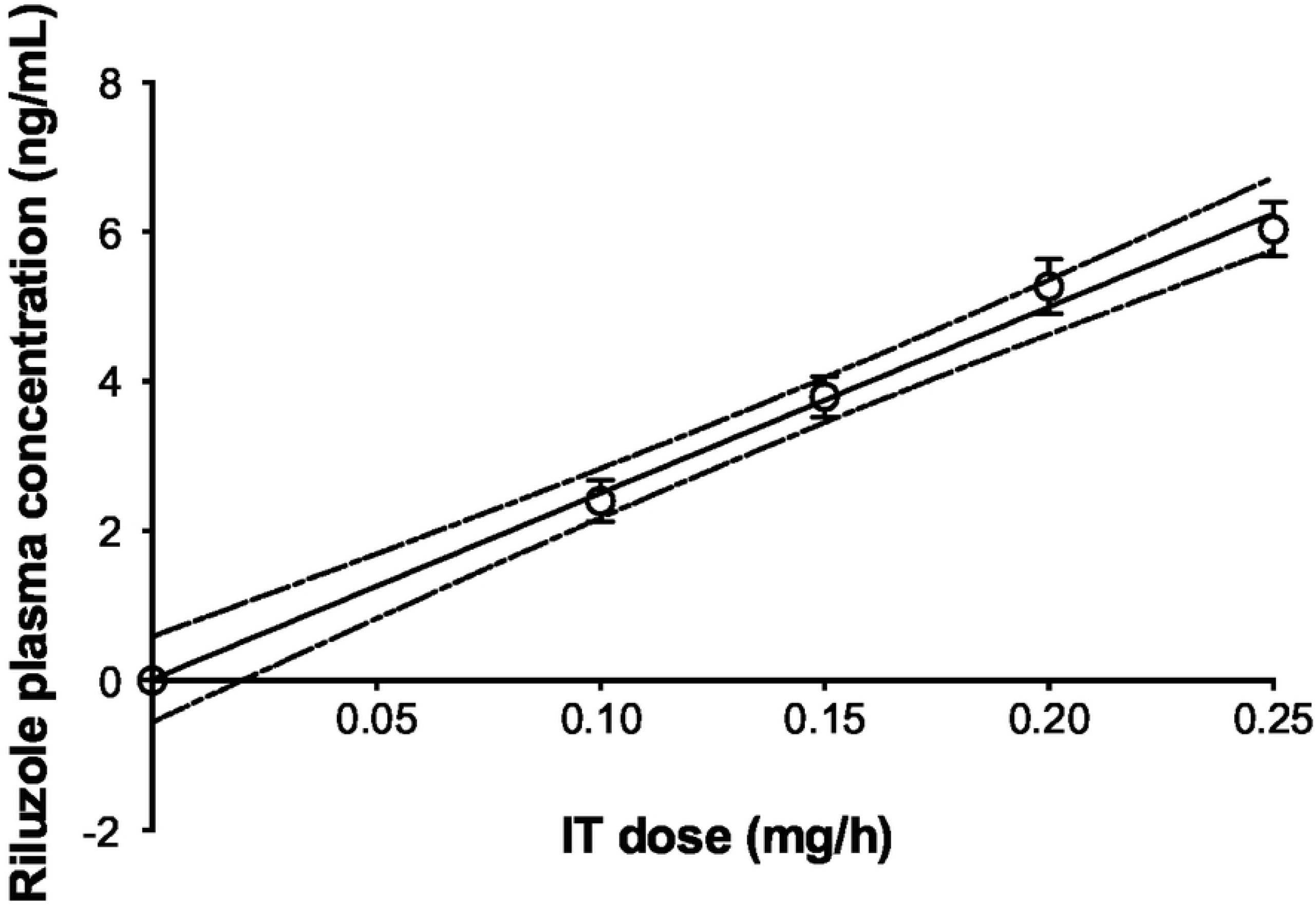
Study 2: Linear regression of riluzole plasma concentration measured during continuous intrathecal infusion at escalating doses of riluzole in dogs. Plasma samples were taken on the fifth day of infusion at each dose, before increasing to the next dose. Regression: Y= 24.91*X + 0.01261 and R^2^= 0.9943. Data shown as means ± SEM

## Results

### Comparison of oral and IT riluzole pharmacokinetics

#### Study 1

Riluzole was detected as soon as 12 hours after IT infusion start, and plasma concentrations rose steadily to reach an apparent steady-state by 48-72 hours (Fig 2). Systemic concentrations of riluzole achieved with IT administration were compared to oral dosing to estimate relative bioavailability (BA). Following a single oral administration of 50 mg of riluzole, the mean C_max_ and AUC_INF_ values for riluzole were 659 ng/mL and 2020 hr*ng/mL, respectively. Following IT infusion of 0.2 mg/hr of riluzole for 148 hours, the mean C_max_ and AUC_0-128hr_, values for riluzole were much lower at 4.15 ng/mL, and 349 hr*ng/mL, respectively. The systemic relative BA with IT infusion (AUC_0-128hr_) was 17.3% of the oral route (AUC_INF_). In addition, the plasma concentrations observed with IT infusion were less variable than the oral route, with CV values ranging from 6.64% to 21.1% for IT vs 72.3% to 163% for oral, indicating greater consistency of plasma concentrations using the infusion system.

### Comparison of Plasma, SC and Brain Concentrations of Riluzole with IT and Oral Delivery

#### Study 2

Infusion of IT riluzole resulted in a linear dose-dependent increase in systemic plasma riluzole concentrations (linear regression, Fig 3). In comparison, concentrations achieved 1 hour after oral riluzole administration were much higher than those achieved with even the highest IT dose (0.2 mg/hr) and much more variable (Fig 4).

**Fig 4.**
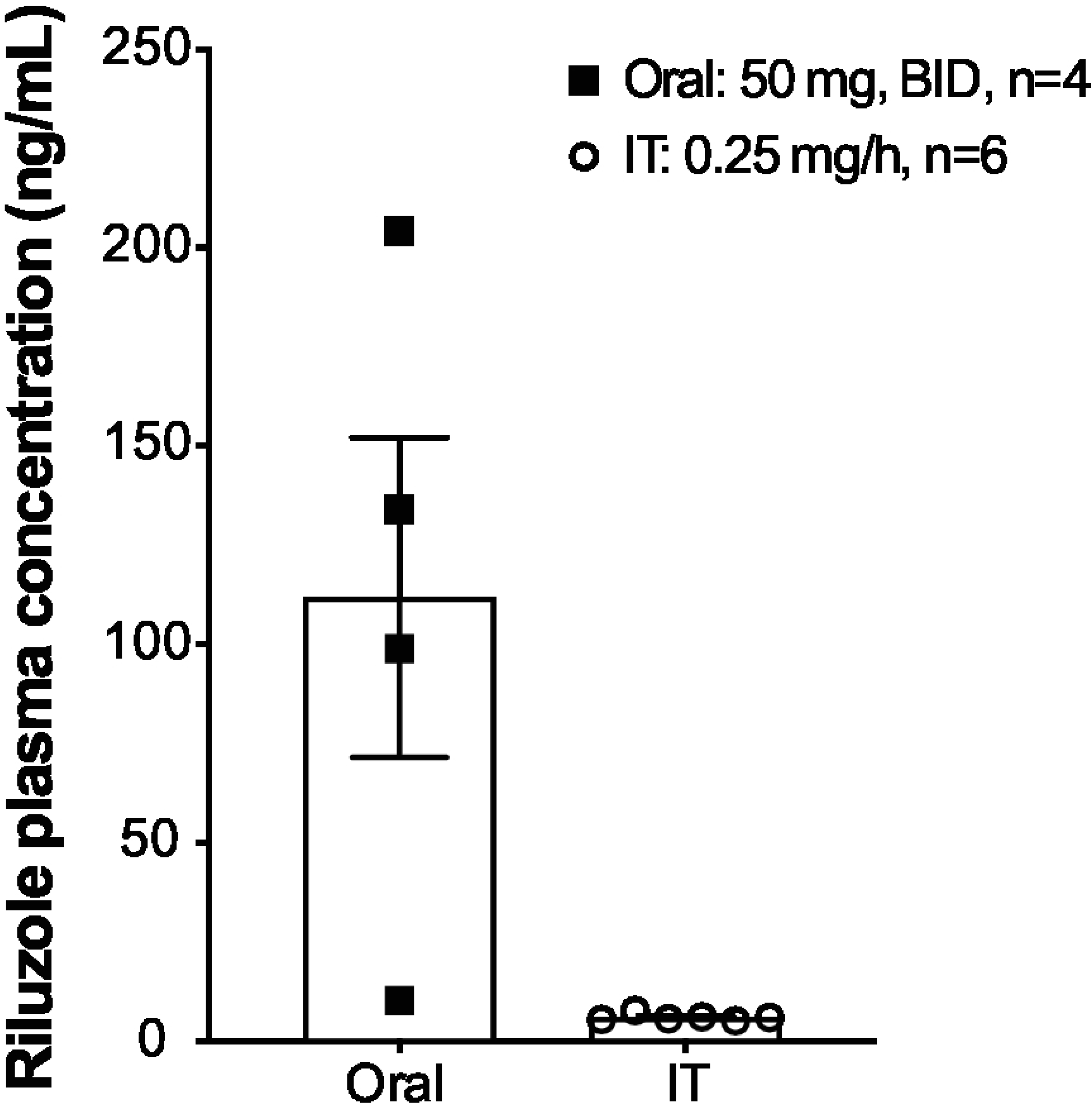
Study 2: Mean plasma concentration of riluzole. Dogs were treated with oral riluzole 50 mg BID for 14 days or IT infusion of riluzole at 0.25 mg/h for 5 days. Plasma samples were taken approximately 60 min after the last oral dose or before stopping IT infusion on day 5. Data shown as means ± SEM

SC and brain concentrations of riluzole were only obtained with the last and highest dose tested (because they required euthanasia for collection), and reflect levels obtained after 5-days of continuous infusion at 0.25 mg/hr. SC data after IT infusion in comparison to those achieved with oral treatment of riluzole one hour after the last dose of a 14-days treatment are shown in Fig 5. With IT infusion, tissue concentrations showed a rostrocaudal gradient with higher levels at the catheter tip and decreased gradually moving caudally from the catheter tip. Drug accumulation rostral to the catheter tip was more limited. This profile of accumulation is similar to what has been observed for other IT drugs [32]. Conversely, within the group of dogs receiving oral riluzole only, the drug concentrations were very consistent along the SC, but variable between dogs. Oral plasma levels significantly correlated with SC tissue concentrations (P<0.05; Fig 6). For most of the sections of the SC examined in the IT group, average concentrations of riluzole were higher than those achieved with 14-days of oral riluzole treatment (at times over a log fold higher). Brain concentrations of riluzole averaged 715 ± 275 ng/g (mean ± SEM; n= 4) with oral riluzole administration and were much lower at 51.0 ± 3.8 ng/g following IT infusion of 0.25 mg/hr (mean ± SEM; n= 6), indicating that IT riluzole infusion at the rate of 50 µl/hr diffuses minimally to the brain.

**Fig 5.**
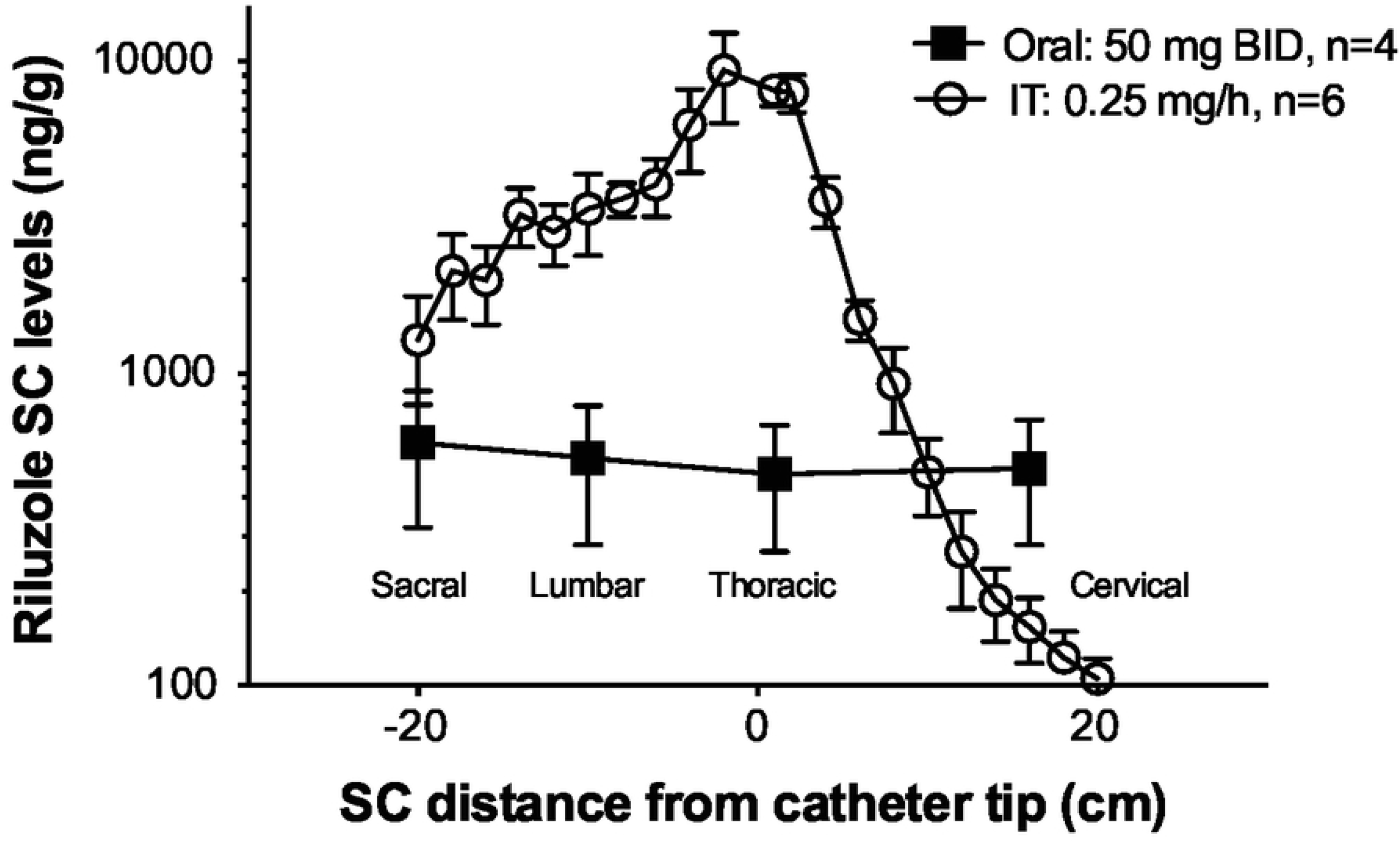
Study 2: SC concentrations of riluzole at terminal necropsy. Dogs were treated for 14 days with oral riluzole (50 mg BID) or continuous IT infusion of riluzole at 0.25 mg/h for 5 days. Tissue samples were taken approximately 60 min after the last oral dose or shortly after stopping IT infusion on day 5. The entire spinal cord was cut in 1 cm segments. The X-axis represents the distance (in cm) of each segment, measured for riluzole content, from the catheter tip location (0) noted at necropsy, or artificially set at T1 segment in non-implanted animals. Approximate anatomical position (sacral to cervical) is indicated. Y-axis is on a Log10 scale. Data shown as means ± SEM

**Fig 6.**
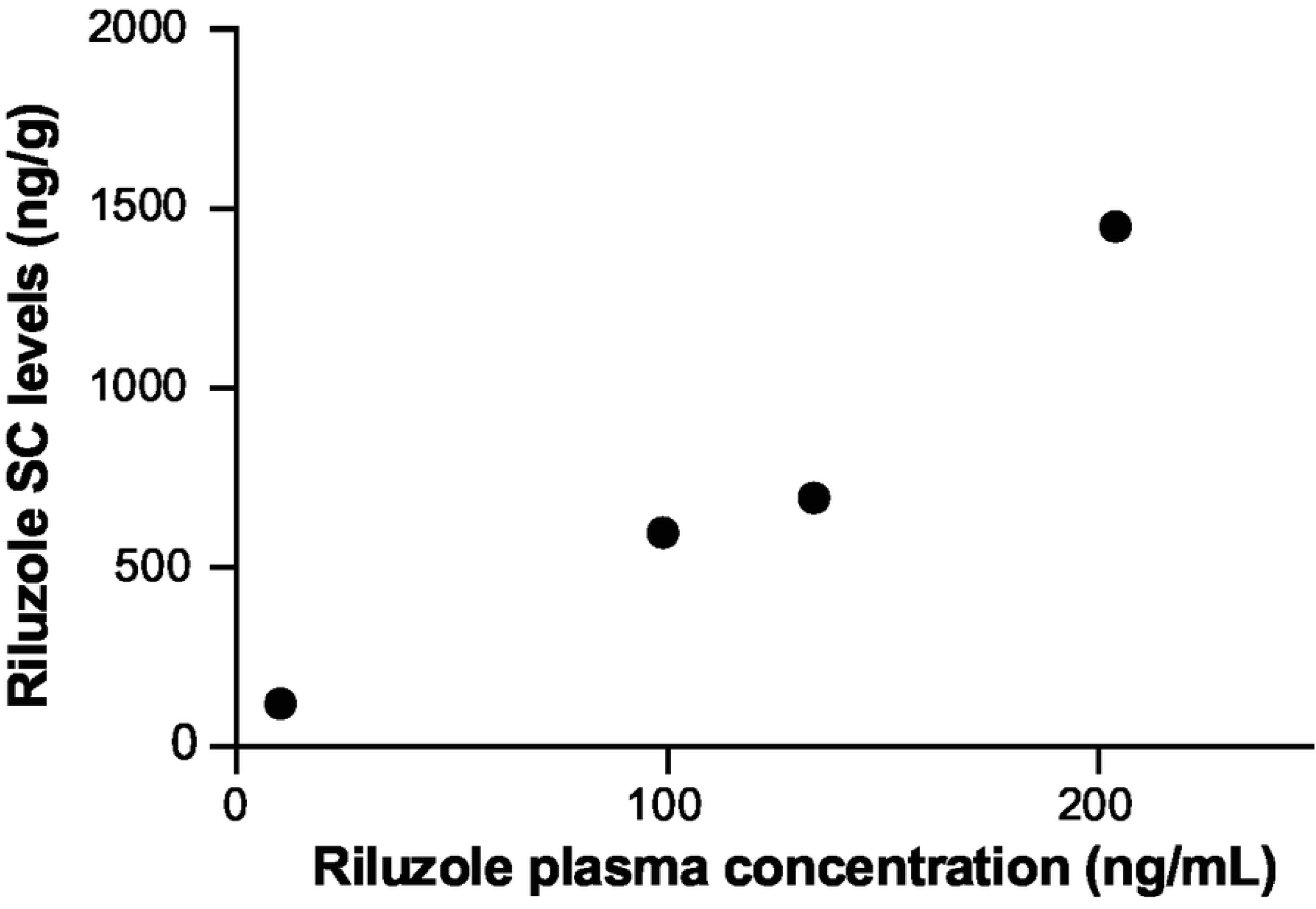
Study 2: Correlation of plasma riluzole to mean riluzole concentrations in the SC. Dogs were treated with oral riluzole at 50 mg BID for 14 days and plasma samples were taken approximately 60 min after the last oral dose. Animals were euthanatized and SC quickly removed thereafter. Data shown as averaged concentration of riluzole in all spinal cord analyzed segments and mean plasma concentrations (R^2^= 0.9435, P= 0.029)

#### Study 3

Three intrathecal doses of riluzole were evaluated in combination with oral riluzole at HED. IT infusion at the dose of 0.1 mg/hr yielded SC concentrations similar to the oral only group while IT infusion of 0.2 mg/hr of riluzole increased SC concentrations of riluzole caudal to the catheter tip above those achieved with oral riluzole alone (Fig 7). Surprisingly, IT infusion at the dose of 0.05 mg/hr yielded SC concentrations below those provided by oral administration, even though the animals also received the oral drug (see discussion). Brain concentrations (ng/g) of riluzole did not differ significantly between groups and were 975 ± 229, 412 ± 183, 827 ± 219 and 653 ± 181 for the groups oral, oral+IT riluzole 0.05 mg/hr, oral+IT riluzole 0.1 mg/hr and oral+IT riluzole 0.2 mg/hr, respectively (mean ± SEM; P> 0.05).

**Fig 7.**
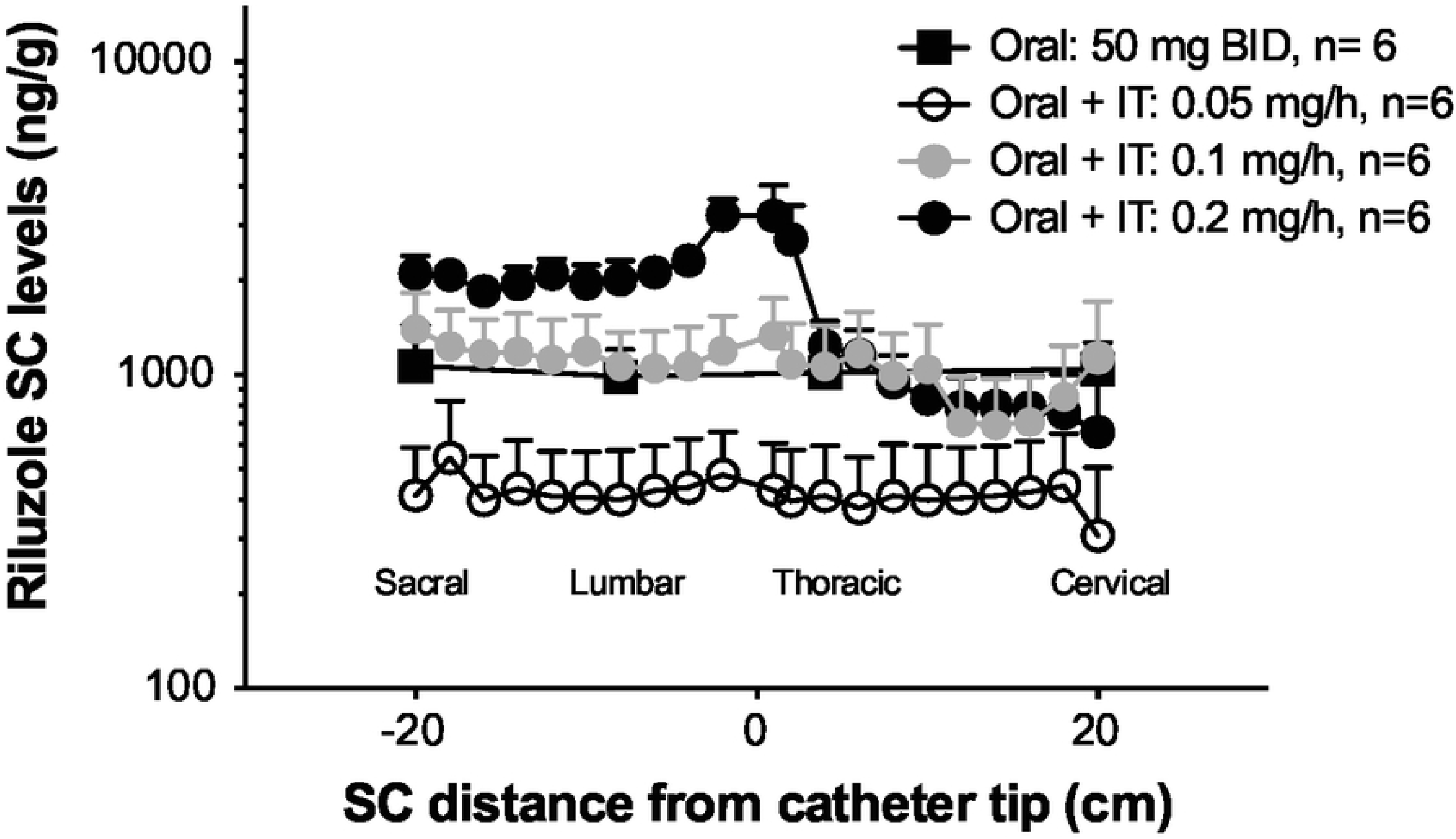
Study 3: SC concentrations of riluzole at terminal necropsy. Dogs were treated for 6-weeks with oral riluzole (50 mg BID) or a combination of oral and continuous IT infusion of riluzole at 0.05 mg/h, 0.1 mg/h or 0.2 mg/h. The entire spinal cord was cut in 1 cm segments. The X-axis represents the distance (in cm) of each segment, measured for riluzole content, from the catheter tip location (0) noted at necropsy, or artificially set at T1 segment in non-implanted animals. Approximate anatomical position (sacral to cervical) is indicated. Y-axis is on a Log10 scale. Data shown as means + SEM

#### Study 4

Three IT doses of riluzole were used in this study in combination with oral treatment at HED. Two doses 0.1 and 0.2 mg/hr bridged back to Study 3 while a higher dose (0.4 mg/hr) extended the dose range previously studied. SC concentrations observed in this study are shown in Fig 8. As in the 6-wk study, IT infusion of riluzole increased SC concentrations above those observed in dogs receiving oral riluzole only. While infusion at 0.1 mg/hr did not increase riluzole concentrations above the oral only group during the 6-week study, this dose elevated SC riluzole above oral drug only after 6 months, with values that were similar to concentrations observed in dogs receiving 0.2 mg/hr. Dogs receiving 0.4 mg/hr did not tolerate this dose as attested by adverse clinical signs (see below). Dogs in this group were euthanized to obtain nervous tissue concentrations at times when adverse clinical signs were observed. Average and maximal SC concentrations in this group were much higher than in those dogs infused with 0.1 or 0.2 mg/hr (note Log10 scale for Y-axis). Brain concentrations (ng/g) of riluzole for oral, oral+IT riluzole 0.1 mg/hr, oral+IT riluzole 0.2 mg/hr and oral+IT riluzole 0.4 mg/hr were 954 ± 202, 1015 ± 296, 845 ± 195 and 1899 ± 153, respectively (mean ± SEM). Brain concentrations of riluzole achieved by the 0.4 mg/hr dose were significantly higher (P<0.05) than achieved for the other groups. This was the first instance of an IT infused dose of riluzole increasing brain concentration beyond those observed with oral drug treatment alone.

**Fig 8.**
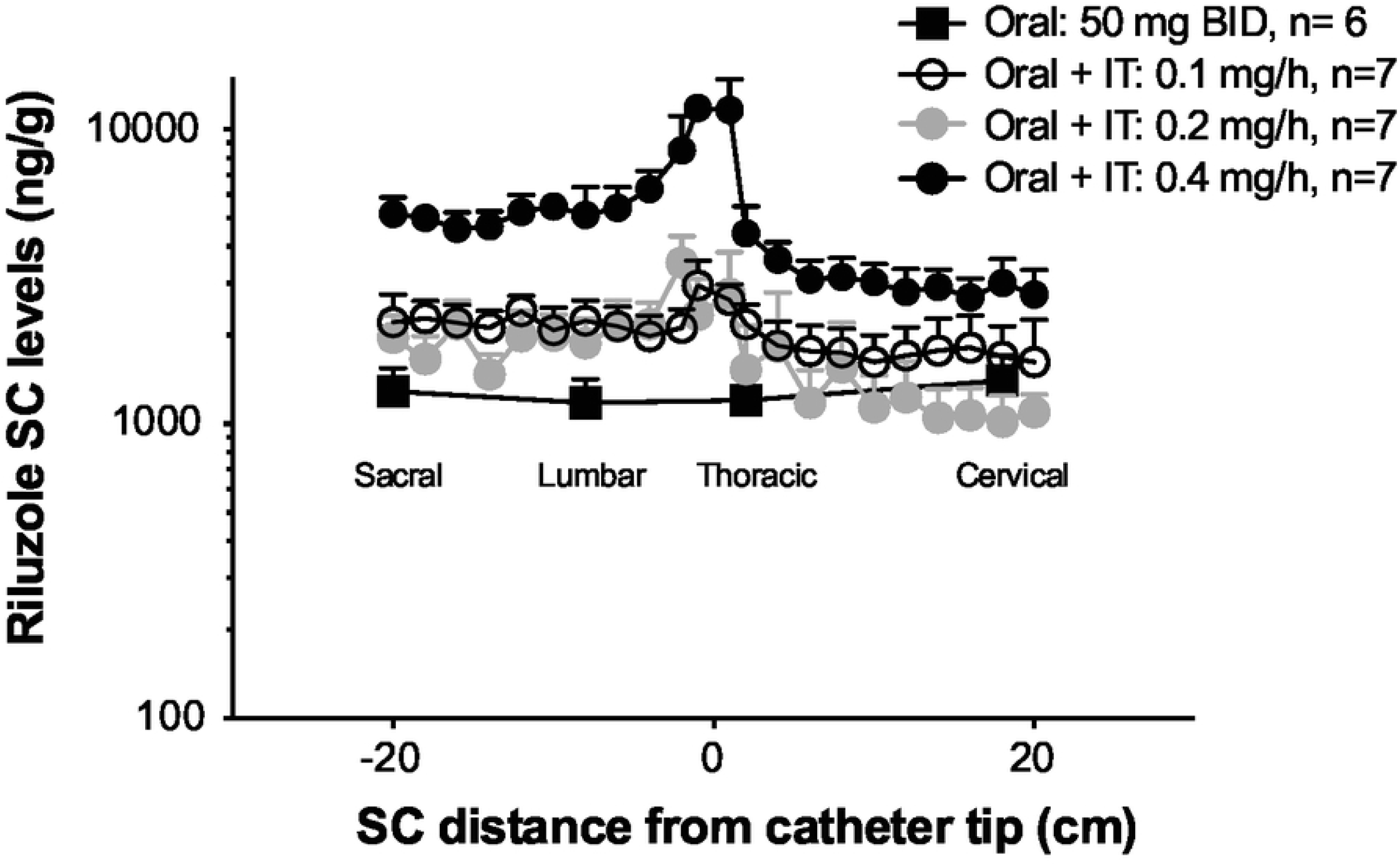
Study 4: SC concentrations of riluzole at terminal necropsy. Dogs were treated for 24-weeks with oral riluzole (50 mg BID) or a combination of oral and continuous IT infusion of riluzole at 0.1 mg/h, 0.2 mg/h or 0.4 mg/h. Dogs receiving IT infusion at 0.4 mg/h were necropsied at various stages of the study due to tolerability issues. The entire spinal cord was cut in 1 cm segments. The X-axis represents the distance (in cm) of each segment, measured for riluzole content, from the catheter tip location (0) noted at necropsy, or artificially set at T1 segment in non-implanted animals. Approximate anatomical position (sacral to cervical) is indicated. Y-axis is on a Log10 scale. Data shown as means + SEM

### Assessments of Tolerability and Safety with IT riluzole

#### Clinical pathology

IT infusion of riluzole was not associated with any significant clinical pathology. Specifically, chronic administration of IT riluzole in combination with oral dosing did not impact hepatic functions as indicated by liver enzymes activities measured in the 6-week (Table 3) and the 6-month Study (Table 4).

**Table 3.** Liver clinical chemistry values at end of Study 3 (Day 42).

|  | *Group 1* | *Group 2* | *Group 3* | *Group 4* | *Group 5* | *Group 6* |
| --- | --- | --- | --- | --- | --- | --- |
| GGT<br>(U/L +/- SD) | 2.7 +/- 0.87 | 2.7 +/- 0.72 | 2.7 +/- 0.23 | 2.5 +/- 0.57 | 3.0 +/- 0.93 | 3.5 +/- 0.85 |
| AST<br>(U/L +/- SD) | 23.2 +/- 4.27 | 26.3 +/- 4.00 | 19.5 +/- 3.73 | 21.7 +/- 4.63 | 24.0 +/- 7.13 | 23.0 +/- 2.91 |
| ALT<br>(U/L +/- SD) | 18.6 +/- 4.28 | 25.6 +/- 14.31 | 20.6 +/- 3.93 | 24.2 +/- 6.58 | 22.0 +/- 2.90 | 21.5 +/- 8.19 |

**Table 4.** Liver clinical chemistry values at end of Study 4 (Day 175).

|  | *Group 1* | *Group 2* | *Group 3* | *Group 4* | *Group 5* | *Group 6* |
| --- | --- | --- | --- | --- | --- | --- |
| GGT<br>(U/L +/- SD) | 3.1 +/- 0.35 | 3/0 +/- 0.53 | 3/0 +/- 0.00 | 3.5 +/- 0.55 | 2.7 +/- 0.76 | 2.5 +/- 0.71 |
| AST<br>(U/L +/- SD) | 23.9 +/- 3.23 | 20.6 +/- 3.74 | 21.8 +/- 2.64 | 24.7 +/- 7.12 | 23.9 +/- 3.98 | 22.0 +/- 1.41 |
| ALT<br>(U/L +/- SD) | 32.5 +/- 3.42 | 25.0 +/- 6.28 | 32.7 +/- 5.09 | 27.8 +/- 7.03 | 31.4 +/- 6.55 | 30.0 +/-<br>12.73 |

#### Clinical Tolerability

Clinical Tolerability was not formally evaluated in Study 1, although no clinical signs were reported. In Studies 2 and 3, all doses of IT riluzole studied were considered to be well tolerated. While occasional mild behavioral deficits were noted throughout both studies, these were not determined to be related to IT infusion of riluzole. The no observed adverse effect level (NOAEL) for IT riluzole in combination with oral dosing at HED was determined to be 0.25 and 0.2 mg/hr for Studies 2 and 3, respectively. In Study 4, the highest IT dose of riluzole (0.4 mg/hr) was not tolerated. All animals in the 0.4 mg/hr group showed moderate to severe clinical signs. Observations included moderate to severe pelvic limb dysfunction, including hind foot paw knuckling, hind limb impairment, hind limb splaying, and/or complete loss of limb function. Behavioral testing scores showed consistently more frequent and severe scores with declines in motor coordination and muscle tone. Neurologic evaluation showed consistent abnormalities in hind limb conscious proprioception and variable withdrawal reflex deficits. Notably, these signs were more pronounced when the observations were made around 1-3 hours after oral administration (estimated plasma C_max_), consistent with an exaggeration of riluzole pharmacological effects. Importantly, these symptoms progressed to a severity that made dosing holidays necessary for all animals, between 14 and 47 days after initiation of continuous IT infusion. After being given a dosing holiday, all dogs in this group recovered from the behavioral deficits induced by 0.4 mg/hr of IT riluzole. However, adverse clinical signs typically returned after roughly the same time period once IT riluzole was re-initiated. Dogs that displayed clinical relapse were then euthanized to establish nervous tissue concentrations of riluzole associated with the clinical signs described above. The dose of 0.2 mg/hr that defined the NOAEL after 6-weeks of continuous infusion in study 2 was equally well tolerated after 6-months of infusion. However, two animals out of 7 animals receiving the low dose of 0.1 mg/hr (previously tolerated in the 6-weeks study) displayed clinical signs including pelvic limb dysfunction, decline in motor coordination, decreased muscle tone, and reduced conscious proprioception. These signs were reversible upon dosing holiday and reappeared in only one of the two animals once treatment was re-initiated. While the higher dose of 0.2 mg/hr was well tolerated, the adverse events in the low dose group precluded the establishment of a NOAEL in this longer study.

#### SC histopathology

In none of the three studies where histopathology was performed did oral administration of riluzole, at approximately the human equivalent dose, produce any histopathological changes in the nervous tissues examined.

In Study 3, histopathologic findings were associated with IT infusion of the THPB-EC vehicle. Representative sections of SC tissue in animals treated with intrathecal saline, vehicle, and riluzole are shown in Fig 9. Compared to intrathecal saline, dosing with intrathecal THPB-EC vehicle was associated with multiple changes including exacerbated catheter track inflammatory reactions, increased spinal cord dorsal and lateral tract degeneration, and increased inflammatory cell infiltrates in the meninges of the brain and SC. This included the presence of a population of vacuolated macrophages, possibly cells that originated from phagocytosis of cell debris or vehicle components. When compared to THPB-EC treated controls, animals treated with a combination of oral and intrathecal infusion of riluzole at all dosing levels showed slight equivocal increases in degeneration of the dorsal and lateral white matter tracts of the SC. None of these histologic changes were considered adverse since they were not associated with any observed alterations in clinical signs.

**Fig 9.**
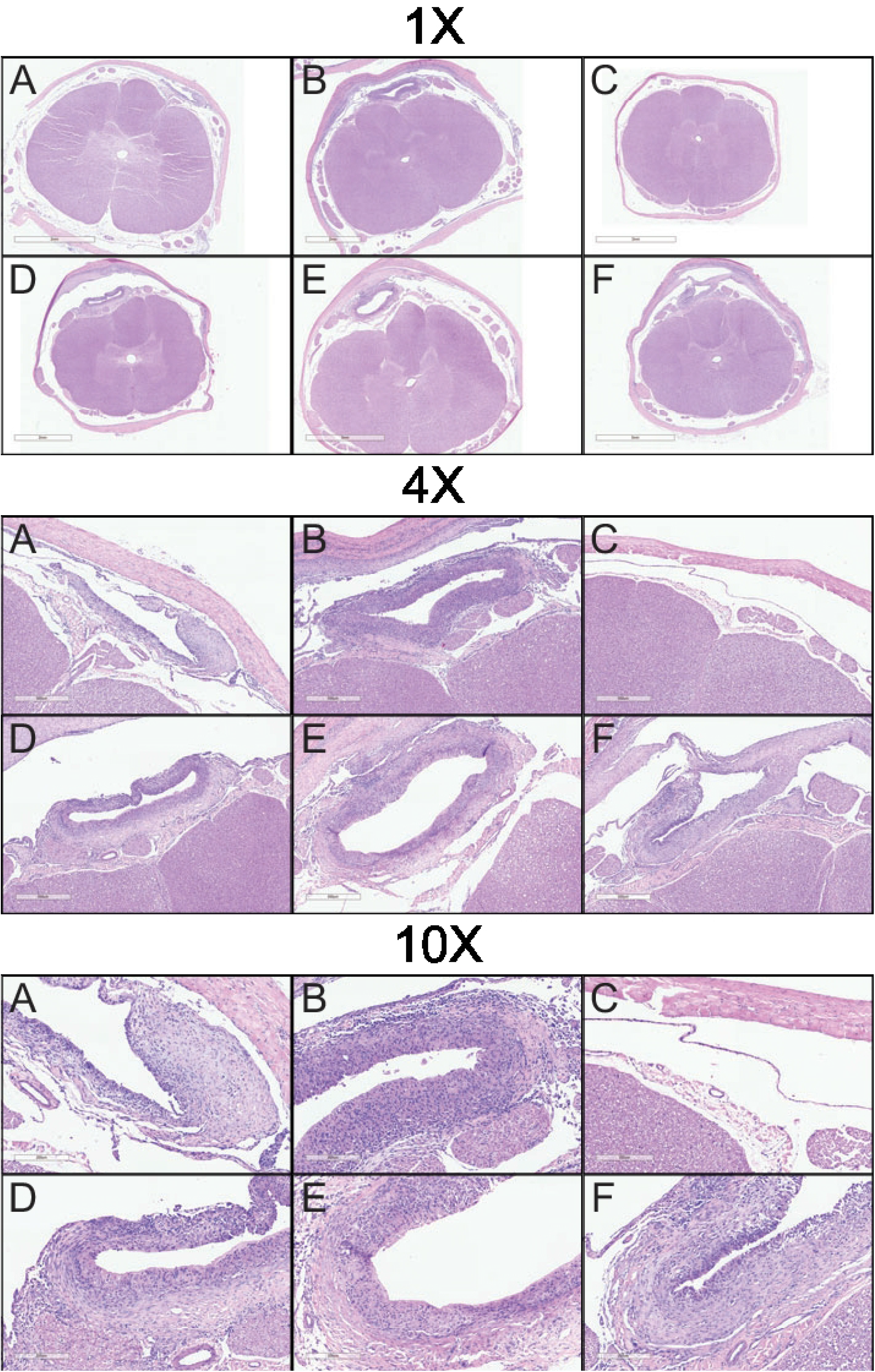
Microscopic images showing representative sections of SC tissue and catheter tracks stained with H&E from animals in Study 3 (6 weeks GLP). Animals were treated with A: IT saline, B: IT vehicle, C: oral riluzole, D: oral + IT riluzole (0.05 mg/h), E: oral + IT riluzole (0.1 mg/h) and F: oral + IT riluzole (0.2 mg/h). Magnification as indicated. The cyclodextrin THPB-EC vehicle was associated with exacerbated catheter track inflammatory reactions as compared to the saline controls. The spinal cord of animals treated with oral riluzole only (C) were histologically within normal limits. Animals treated with IT riluzole showed equivocal, slight increases in SC degeneration (not visible here) compared to vehicle controls

Study 4 revealed histopathological findings in animals dosed with 0.1 mg/h or 0.2 mg/h that were similar to those observed in Study 3 (Fig 10). As in Study 3, the Vehicle 4% THPB-EC IT caused increased tissue reactions compared to saline IT. These included exacerbated catheter track inflammatory reactions, increased spinal cord dorsal and ventral tract degeneration, and increased inflammatory cell infiltrates in the brain and SC meninges and spinal nerve root epineurium (contiguous with meninges) including vacuolated macrophages. In this study, minimal spinocerebellar tract degeneration attributed to spinal cord changes was also seen. Effects of riluzole in animals treated with 0.1 or 0.2 mg/hr were similar to those seen in animals with the same doses from Study 3, with the exception of the one animal in the 0.1 mg/h dose group that developed clinical signs upon treatment re-initiation. These effects included increases in degeneration of the dorsal tracts white matter of the spinal cord as well as slight increases in degeneration of the spinocerebellar tracts. In animals treated with 0.4 mg/h, riluzole caused additional exacerbations of catheter track inflammatory reactions and inflammatory changes in the parenchyma of the spinal cord. No riluzole effects were seen in the spinal nerve roots, DRG, or peripheral nerves. In some animals with clinical signs, the relatively low intensity of changes seen in the SC microscopically would not be expected to induce the clinical signs described. This suggested the possibility that some of the clinical signs could be due to a functional mechanism that are not evident morphologically via microscopic examination (e.g. exaggerated drug pharmacology).

**Fig 10.**
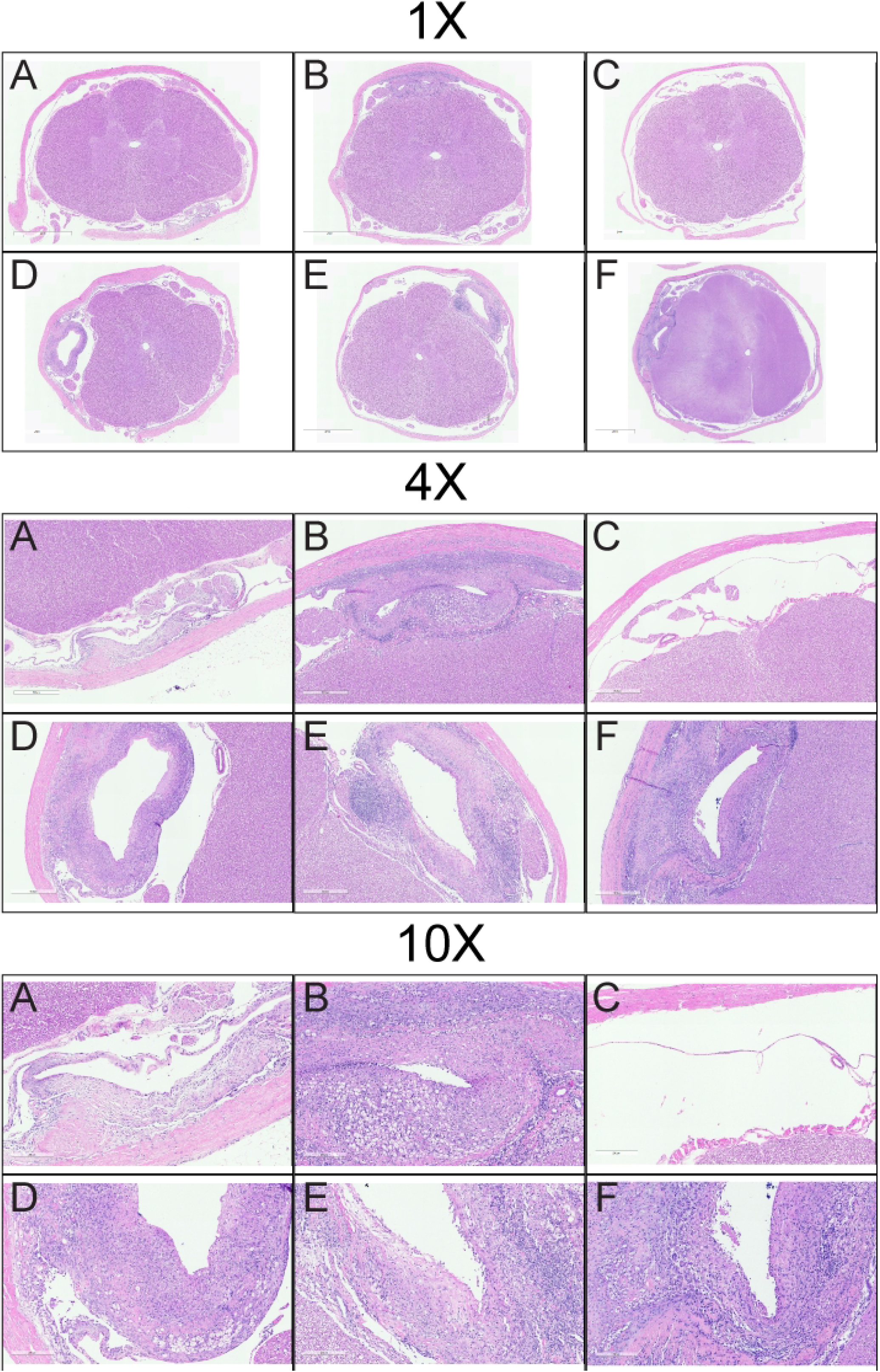
Microscopic images showing representative sections of SC tissue and catheter tracks stained with H&E from animals in Study 4 (6 month GLP). Animals were treated with A: IT saline, B: IT vehicle, C: oral riluzole, D: oral + IT riluzole (0.1 mg/h), E: oral + IT riluzole (0.2 mg/h) and F: oral + IT riluzole (0.4 mg/h). Magnification as indicated. The cyclodextrin THPB-EC vehicle was associated with exacerbated catheter track inflammatory reactions as compared to saline controls. The spinal cord of animals treated with oral riluzole only (C) were histologically within normal limits. Animals treated with 0.1 or 0.2 mg/h IT riluzole showed increases in SC degeneration (not visible here) compared to vehicle controls. Animals treated with 0.4 mg/h IT riluzole sowed additional exacerbations of catheter track inflammatory reactions and inflammatory changes in the parenchyma of the spinal cord (visible here)

## Discussion

More than 20 years after the initial demonstration that riluzole could benefit patient with ALS, minimal effort has been made to improve its therapeutic potential. Mainly, dexpramipexole (a benzothiazole derivative of riluzole) was developed but failed to display efficacy in the clinics [33]. Additionally, to facilitate administration in patients with ALS driven swallowing difficulties, an oral suspension was approved (Tiglutik®), and a sublingual form is currently under development (Biohaven Pharmaceuticals). We aimed to develop, characterize, and evaluate the safety of IT delivery of riluzole.

Lipophilic drugs tend to show discrete patterns when infused in the IT space, including reduced retention, diffusion to lipid compartments, and rapid plasma clearance [34]. In the case of riluzole, our data in dogs showed a very favorable profile when infused in the IT space. IT infusion yielded minimal but detectable plasmatic concentrations of riluzole while enabling retention of the drug in the SC. At the low doses that we administered to the CSF, the systemic bioavailability (BA) of IT riluzole was low when compared to that of the oral route. Therefore, when administered as a combination treatment, IT riluzole did not increase further the plasma levels provided by oral riluzole, even at the highest IT dose tested. Finally, the linear dose-response curve established in our dose escalation study provided evidence of the robustness and reliability of the delivery system even at the low infusion rate of 50 µl/hr (1.2 ml/day).

As could be expected from the plasma data, IT infusion did not impact the dog liver. Measures of hepatic function, including transaminases levels, remained unchanged throughout our studies (Table 3 and 4). Thus, it was possible to increase on-target delivery of riluzole to the spinal cord without concurrently raising the risk of further elevating liver enzymes. Further, in the current study, bioanalytical analysis of riluzole showed that IT infusion alone provided minimal concentrations of the drug in the brain (albeit a potential downside if riluzole has a therapeutic benefit at this level), and did not increase the brain levels provided by oral administration when given as a combination treatment. The only instance of IT infusion of riluzole increasing brain concentration above those observed with oral treatment alone was found in the 0.4 mg/hr group in Study 4. This may have been due to the higher infusion rate used in this group or saturation of drug efflux transporters at the blood brain barrier (see below). This finding, in combination with lower systemic levels, could also limit other side-effects of riluzole, including asthenia and digestive system issues (nausea, anorexia). Reducing these effects, which were known to lead to patient withdrawal in the dose-range finding clinical studies, might help with patient compliance. [14, 18].

In addition to the side-effects of oral riluzole, there is also a very high inter-individual variability of serum concentration, which in turn means there is high inter-individual on-target (e.g. CNS) concentrations. As such, there is certainly a subset of ALS patients in whom riluzole concentrations achieved with oral dosage are likely too low to achieve full efficacy [35]. The same variability in oral BA was seen in dogs receiving oral riluzole, as indicated by the plasma levels measured in Study 2. Importantly, in these animals, brain and SC tissue levels correlated with riluzole plasma levels. Since IT infusion diffuses minimally to the brain and plasma, this indicates that the brain riluzole concentrations observed are almost exclusively driven by the plasma levels provided by only the oral dosing. In Study 3, despite receiving oral riluzole at the same dosage, the SC levels in the IT low dose group were significantly lower than in the oral group. However, brain concentrations in the IT low dose group were also lower than the oral only group. This indicates that plasma concentrations were likely reduced in these animals, and that the observed spinal concentrations were mostly the result of IT infusion. Similarly, inter-animal variability in plasma concentration likely explained the lack of difference in SC concentrations between the 0.1 mg/hr and 0.2 mg/hr groups seen in Study 4, which did not replicate Study 3 findings. Overall the findings here confirm that oral riluzole is a highly variable therapeutic, and that IT riluzole has the potential to provide more consistent, though spatially limited concentrations of riluzole.

Importantly, SC concentrations measured in our studies represent tissue levels that are sustained throughout treatment as a result of continuous IT infusion. Thus, IT infusion provides a more homogenous administration of the drug, likely affording more constant neuroprotection to the targeted SC motor neurons. Our studies revealed that IT infusion alone or in combination with oral administration significantly increased the SC concentrations of riluzole. Notably, results in IT treated animals were compared to control oral groups where the tissues were collected near C_max_ of oral administration. Thus, it is likely that our data underestimate the BA of IT riluzole in the SC, as the measured riluzole levels are maintained continuously with this route. IT infusion resulted in a gradient of riluzole concentrations in the SC tissue, with higher concentrations at the catheter tip, a steep concentration decrease rostrally, and a more gradual concentration decrease caudally. This concentration profile is very similar to other drugs delivered by the IT route [32, 36], and could be exploited to develop targeted segmental approaches, for example by aiming at providing higher concentration of riluzole to the SC segment responsible for diaphragmatic functions.

Evaluation of IT riluzole in patient would require an evaluation of the safety profile of the treatment, however, given that oral riluzole is standard of care it would be both unethical and unfeasible to evaluate IT riluzole in isolation. Therefore, we assessed the safety/toxicity of IT riluzole infusion in combination with oral administration at human equivalent dose in two consecutive GLP studies. The IT dose of 0.4 mg/hr was not tolerated and defined the maximum tolerated dose of the IT riluzole/oral combination. The adverse clinical signs observed at this dose aligned with an exaggeration of the pharmacological profile of riluzole. Importantly, riluzole concentrations in the SC of these animals reached levels similar to those previously reported as defining the upper limit of tolerability of oral riluzole at C_max_ in the rat SC (described in the SBA for Rilutek® NDA 20-599 [51]). Continuous IT infusion at the dose of 0.2 mg/hr for up to 6-month was well tolerated and was not accompanied by any clinical signs or significant clinical pathology. However, two animals at the lower dose of 0.1 mg /hr experienced hindlimb impairments that required treatment holiday in the 6-month study, but not in the 6-weeks study, which prevented the establishment of a NOAEL in the more extended study. We were not able to find any explanation for this discrepancy, suggesting that there is an individualized response to IT riluzole treatment. It is also possible that this difference was caused by a peculiar anatomical positioning of the catheter, which could not be identified at necropsy. Indeed, catheters underwent significant movement throughout our studies (as indicated by the changes in position from the time of placement to the time of necropsy), and a catheter displacement near a nerve root could have been responsible for these adverse clinical signs. Alternatively, CSF flow dynamics are not uniform, and the catheter tip may have been transiently positioned in a pocket of stagnant CSF, yielding artificially elevated levels of riluzole in the SC or a nerve root. Further studies on implantation, catheter position (acute and chronically), and catheter use will be necessary to determine if this is a causal factor. Importantly, the clinical signs observed in our studies were fully reversible upon treatment cessation. Altogether our clinical data suggest that, except for complications associated with the IT catheter, IT administration of the riluzole formulation was well tolerated in the dog at doses up to 0.2 mg/hr.

Histopathological assessment of SC tissue revealed multiple changes in all animals that received IT infusion, including saline-treated animals. This illustrates a challenge of the IT route of administration since the implantation of IT catheters may alone induce multiple histological changes [37]. It is thus essential to note that our 6-month study, while tolerated in most animals, was considerably longer than most safety studies for intrathecally delivered test articles [38-40]. Some of the changes observed in that study may have been due in part to this long duration leading to exacerbated tissue reactions. In addition, we designed our studies as a platform to support the development of IT riluzole in humans, which required the use of a human device. Hound dogs were chosen as they best enable the subcutaneous implantation of the large 40 ml pump from Medtronic. The hound dog SC anatomy and larger diameter spinal canal was also a better fit for the implantation of the Ascenda catheter. Nonetheless, the spinal canal diameter in large dogs measures about 0.85 cm with a SC diameter of around 0.5 cm [41], while the Ascenda catheter diameter is 0.12 cm. This indicates that there was still very little space to accommodate the human catheter, which could have impacted responses to the experimental system in dogs. Since the human spinal canal diameter measures 1.5-3.2 cm with a SC diameter of around 0.7-1.4 cm [42, 43], placement and maintenance of an IT catheter in humans is less challenging and fraught with fewer reactions. In animals treated with 0.1 and 0.2 mg/h, the majority of histologic changes seen in treated animals as compared to saline controls were attributed to the THPB-EC vehicle, suggesting that development of a new vehicle could be necessary for later stage development of the therapy. In addition to altering the safety profile of IT riluzole, these increased tissue reactions may have modified the biodistribution profile of the drug, in particular in the 6-months GLP study. It is important to note that the severity of these reactions is within the range observed in similar studies of other standard of care IT therapies for nonterminal diseases, which have been administered in humans for many years, including opiates, baclofen, bupivacaine and ziconotide [44]. Consent for these therapies includes the rare possibility that patients will develop catheter tip granulomas that may themselves require therapy and preclude further IT treatment [45, 46].

Overall, these studies establish the development, characterization and safety profile for IT delivery of riluzole and he next steps will involve characterizing is therapeutic efficacy. In addition to higher and more consistent dosing of riluzole on target, we believe this delivery method offers another advantage. Recent evidence indicates that riluzole may not reach sufficient CNS exposure for reasons beyond what we have discussed. There is a known progressive, disease-driven upregulation of the drug efflux transporter P-glycoprotein (P-gp) and breast cancer resistance protein (BCRP) at the blood brain barrier. The activity of both of these cellular pathways progressively increases in the SC of both the SOD1 mouse model and in ALS patients [47]. Riluzole is a substrate for the p-glycoprotein system, which limits its ability to penetrate the blood brain barrier. Accordingly, administration of riluzole in combination with the P-gp/BCRP inhibitor elacridar in SOD1 mice increased riluzole concentration in the CNS and extended the survival benefit provided by riluzole alone [20]. Consistent with the concept that progressively increased efflux of riluzole from the SC hinders riluzole efficacy, clinical observations indicated that riluzole was far more effective in the first 12 months of treatment than in the following year [13, 48]. Continuous IT infusion of riluzole at tolerated doses up to 0.2 mg/hr can significantly elevate SC levels 2 to 3 times above those provided by oral riluzole and, in mice lacking the P-gp transporters, riluzole concentrations in the CNS only doubled [20]. It is thus likely that IT administration would be able to overcome this disease driven decrease in SC retention of the drug. Intriguingly, a recent retrospective study indicated that riluzole prolongs survival in the very last clinical stage of ALS when the IT route could be best suited to maintain appropriate riluzole levels over oral dosing [49]. However, it should be noted that these studies did not parse the benefits of increased riluzole in the brain or in the spinal cord, or both. Therefore, if the effect of riluzole is only beneficial on the motor neurons in the brain/brainstem, the IT delivery of the riluzole would not be effective.

In conclusion, our data show that IT infusion of riluzole provides a consistent and elevated SC concentration without increasing the brain or systemic levels. When adverse clinical signs effects emerged, they were associated with the higher SC riluzole concentration in the affected animals, and their nature suggested an exaggeration of the drug pharmacological profile. Supporting this, in some animals, changes seen on histopathologic examination were not present to the extent that would be expected to be associated with the adverse clinical signs observed. Importantly, these adverse clinical signs could be detected early before worsening and were reversible upon cessation of treatment. While our experimental data suggest a narrow therapeutic index for the combination of oral and IT riluzole, the anatomical differences between dog and humans spinal canal suggest that, with careful consideration of the starting dose and close monitoring, IT infusion at doses up to 0.2 mg/hr would be tolerated in humans. We believe that using IT administration to create increase riluzole concentrations in the SC and improve the consistency of delivery within individual patients could increase the drug efficacy for the treatment of ALS, but this will need to be empirically determined in future studies. To that intent, a multiple ascending dose escalation clinical trials is being planned to evaluate the safety of IT riluzole in patients with ALS.

